# The Molecular Determinants of a Universal Prion Acceptor

**DOI:** 10.1101/2024.03.01.582976

**Authors:** Hamza Arshad, Zeel Patel, Zaid A.M. Al-Azzawi, Leyao Li, Genki Amano, Surabhi Mehra, Shehab Eid, Gerold Schmitt-Ulms, Joel C. Watts

**Author notes:** To whom correspondence should be addressed at: Krembil Discovery Tower, Rm. 4KD481, 60 Leonard Ave., Toronto, ON, Canada, M5T 0S8.

## Abstract

In prion diseases, the species barrier limits the transmission of prions from one species to another. However, cross-species prion transmission is remarkably efficient in bank voles, and this phenomenon can be recapitulated in mice by expression of the bank vole prion protein (BVPrP). The molecular determinants of BVPrP’s ability to function as a universal or near-universal acceptor for prions remain incompletely defined. Building on our finding that cultured cells expressing BVPrP can replicate both mouse and hamster prion strains, we conducted a systematic analysis to identify key residues in BVPrP that permit cross-species prion replication. Consistent with previous findings, we demonstrate that residues N155 and N170 of BVPrP, which are absent in mouse PrP but present in hamster PrP, are critical for cross-species prion replication. Additionally, BVPrP residues V112, I139, and M205, which are absent in hamster PrP but present in mouse PrP, are also required to enable replication of both mouse and hamster prions. Unexpectedly, we found that residues E227 and S230 near the C-terminus of BVPrP severely restrict the accumulation of prions following cross-species prion challenge, suggesting that they may have evolved to counteract the inherent propensity of BVPrP to misfold. PrP variants with an enhanced ability to replicate both mouse and hamster prions displayed accelerated spontaneous aggregation kinetics *in vitro*. These findings suggest that BVPrP’s unusual properties are governed by a key set of amino acids and that the enhanced misfolding propensity of BVPrP may enable cross-species prion replication.

## Introduction

Prions are infectious protein aggregates that give rise to animal and human neurodegenerative diseases collectively referred to as prion disorders (1, 2). Naturally occurring prion diseases include scrapie in sheep, chronic wasting disease in deer and elk, and Creutzfeldt-Jakob Disease in humans. Prions are composed of a glycosylphosphatidylinositol (GPI)-anchored protein, the prion protein (PrP), which is encoded by the *Prnp* gene and is expressed within cells of the central nervous system (3). PrP can exist in two conformational states: cellular PrP (PrP^C^), which possesses an intrinsically disordered N-terminal domain and an α-helical C-terminal domain, and a disease-causing conformer termed PrP^Sc^, which forms aggregates and is composed almost entirely of β-sheets (4–6). Distinct PrP^Sc^ structures give rise to different prion strains that have unique pathogenic properties (7, 8). PrP^Sc^ can template the conversion of PrP^C^ into additional copies of PrP^Sc^, thus resulting in the accumulation and spread of prions within the brain (9). While prion diseases can occur sporadically due to spontaneous generation of PrP^Sc^, the ability of PrP^Sc^ to self-propagate means they can also occur via an infectious aetiology. However, a species or transmission barrier limits the ability of prions to pass from one species to another. These barriers are dictated by amino acid sequence differences between PrP^C^ and PrP^Sc^ as well as the structural compatibility of PrP^C^ for a given prion strain (10, 11). When PrP^C^ and PrP^Sc^ are derived from the same species, and therefore have identical amino acid sequences, efficient templating occurs, leading to robust disease transmission. During cross-species prion transmission, differences in amino acid sequence between PrP^C^ expressed by the host and PrP^Sc^ can prevent or impede disease transmission and can lead to prion strain mutation (12–14).

Bank voles (*Myodes glareolus*) present an exception to the species barrier, as they can be efficiently infected with prion strains from a wide range of animals with at least partial maintenance of prion strain properties (15–22). Transgenic mice that express bank vole (BV) PrP are similarly susceptible to cross-species prion infection, arguing that the sequence of BVPrP is structurally compatible with PrP^Sc^ from many different species (23, 24). Moreover, BVPrP functions as a permissive substrate for cross-species prion replication in several *in vitro* prion replication paradigms (25–27), expression of wild-type or mutant BVPrP containing isoleucine at polymorphic codon 109 in mice can lead to spontaneous prion formation (28–31), and recombinant BVPrP can be readily polymerized into infectious aggregates (32–34). Collectively, this suggests that BVPrP is inherently prone to adopting misfolded conformations, potentially allowing it to serve as a universal or near-universal substrate for prion replication with relaxed sequence identity requirements.

The molecular determinants that govern the anomalous behavior of BVPrP in prion transmission experiments remain incompletely defined. As neither mouse (Mo) nor hamster (Ha) PrP enable cross-species prion transmission, the atypical properties of BVPrP must be enciphered within the 7-8 amino acid sequence differences that exist between these three proteins. Previous studies have implicated subsets of BVPrP-specific residues in facilitating cross-species prion replication. For instance, the presence of specific asparagine residues in BVPrP may facilitate prion replication via formation of steric zippers in the final or intermediary structure of PrP^Sc^ (35). Indeed, when PrP sequences between various species of mice and voles were compared, the presence of asparagine at residues 155 and 170 were inferred to be critical for cross-species prion conversion (17). More recently, the importance of residues E227 and S230 in BVPrP, which are in the C-terminal region near the GPI anchor attachment site, has been highlighted. These two residues have been suggested to function as a “linchpin” domain within BVPrP, in that they gatekeep the conversion of BVPrP^C^ into PrP^Sc^ by several different prion strains (36). Additionally, transgenic mice that overexpress MoPrP containing these two BVPrP C-terminal residues develop a spontaneous illness that displays certain neuropathological signs of prion disease (37). However, a comprehensive analysis of the key amino acids that enable BVPrP to function as a universal prion acceptor remains lacking.

Genetically modified mice and *in vitro* prion replication systems have been widely employed to discern the PrP residues that govern cross-species prion replication. However, both paradigms suffer from important drawbacks. For instance, the generation of transgenic or knock-in mice requires significant financial and time commitments, whereas *in vitro* systems risk being unphysiological as prion replication occurs in a cell-free environment. Cultured cell models, on the other hand, represent a more tractable system for studying prion replication as they provide the benefit of using intact cells that express membrane anchored PrP^C^ with its correct post-translational modifications (38). CRISPR/Cas9 gene editing has expanded the range of prion strains that can be studied in cultured cells (39). For instance, CAD5 cells, which can be infected with several different strains of mouse prions (40, 41), can be ablated for endogenous MoPrP (CAD5-PrP^-/-^ cells) and then used as a blank canvass to express PrPs from different species (42, 43). Recently, we demonstrated that CAD5-PrP^-/-^ cells expressing BVPrP enable cross-species prion replication following challenge with several different strains of mouse or hamster prions (44). In this study, we utilized CAD5-PrP^-/-^ cells expressing mutant or chimeric PrPs to systematically identify the amino acid determinants that permit BVPrP to enable cross-species prion replication.

## Materials and Methods

### Cell culture

CAD5-PrP^-/-^ cells were generated by CRISPR/Cas9 gene editing of the mouse catecholaminergic cell line CAD5, which is a subclone of the original Cath.a-differentiated (CAD) line (40, 42, 45). All experiments utilized CAD5-PrP^-/-^ clone D6 (42). Cells were cultured in growth medium, which consists of Opti-MEM media (ThermoFisher Scientific #31985088) supplemented with 6.5% (v/v) fetal bovine serum (ThermoFisher Scientific #12483020) and 1X GlutaMAX (ThermoFisher Scientific #35050061). Cells were cultured at 37 °C in a 5% CO_2_ humidified incubator and were maintained by routine passaging at a dilution of 1:5 or 1:3 every 3-4 days. To passage the cells, they were first washed with PBS (ThermoFisher Scientific #14190144) and then treated with enzyme-free cell dissociation reagent (Millipore Sigma #S-014-B). The cells were incubated for 1-3 minutes in a 37 °C incubator during the dissociation step and then plated in fresh growth medium. Stably transfected polyclonal CAD5-PrP^-/-^ lines were cultured in growth medium containing 0.2 mg/mL G418 (BioShop #GEN418.5). Monoclonal lines of stably transfected CAD5-PrP^-/-^ cells were cultured in growth medium lacking G418 as previously described (44).

### Generation of polyclonal stable cell lines

Expression plasmids encoding wild-type bank vole PrP (M109 variant), mouse PrP, or hamster PrP were generated by cloning the respective *Prnp* open reading frames between the *BamHI* and *Xbal* sites of the vector pcDNA3 (42, 44). The MoC1, MoC2, and HaC1 chimeras were generated by gene synthesis (BioBasic Canada). Other chimeric- and mutant PrP-expressing plasmids were generated by site-directed mutagenesis. For generation of stably transfected CAD5-PrP^-/-^ lines, cells were plated at a density of 700,000 cells/well in 6-well plates, and then transfected with a plasmid encoding the desired gene. For transfection, 2 µg of plasmid DNA was mixed with 4 µL of Lipofectamine 2000 (ThermoFisher Scientific #11668019) in 300 µL of Opti-MEM without any additives. This mixture was added to 1.5 mL of Opti-MEM and then applied to CAD5-PrP^-/-^ cells for 24 h. Next, the cells were washed with PBS and detached using tenzyme-free dissociation reagent. The entirety of the transfected well was then transferred into a 10-cm dish and cultured in medium containing 1 mg/mL G418. The cells were selected over a period of 12-14 days and then expanded. No clonal selection was performed.

### Cell Lysis and immunoblotting

Cells were washed with cold PBS, and then incubated with lysis buffer (50 mM Tris-HCl pH 8.0, 150 mM NaCl, 0.5% (w/v) sodium deoxycholate, and 0.5% (v/v) NP-40) for 30-60 s, swirled, and then lysates collected. A cell scraper was occasionally used when lysing with smaller volumes. Lysates were incubated on ice for 15 min with intermittent vortexing (30 s), and then centrifuged at 5,000x *g* for 5 min to pellet debris. Protein concentrations in cell lysates were determined using the bicinchoninic acid (BCA) assay (ThermoFisher Scientific #23227). For analysis of PrP^C^ levels, protein was prepared for immunoblotting in 1X Bolt LDS sample buffer (ThermoFisher Scientific #B0007) and then heated for 10 min at 70 °C. The samples were run on 10% Bolt Bis-Tris Plus Mini gels (ThermoFisher Scientific #NW00100BOX for 10-well gels) for 35 min at 165 V. Transfer to Immobilon-P PVDF membranes (Millipore Sigma #IPVH00010) was performed for 1 h at 25 V in Tris-glycine transfer buffer (100 mM Tris-HCl pH 8, 137 mM glycine). Membranes were blocked for 1 h at 22 °C in blocking buffer (5% (w/v) skim milk in TBS containing 0.05% (v/v) Tween-20 (TBST)), and then incubated overnight at 4 °C with antibodies diluted in blocking buffer. The following anti-PrP antibodies were used: recombinant humanized Fab HuM-D13 (1:10,000), recombinant humanized Fab HuM-P (1:10,000), and mouse monoclonal POM1 (1:5,000; Millipore Sigma #MABN2285) (46–48). The HuM-D13 antibody was provided by Stanley Prusiner (University of California San Francisco). The following day, membranes were washed 3X with TBST (5-10 min per wash), and then incubated with horseradish peroxidase-conjugated secondary antibodies (Bio-Rad #172-1011 or 172-1019, or ThermoFisher Scientific #31414) diluted in blocking buffer for 1 h at 22 °C. The membranes were again washed 3X with TBST (10 min per wash). The blots were developed using Western Lightning ECL Pro (Revvity #NEL122001EA) and then chemiluminescent signal was captured by exposure to HyBlot CL x-ray film (Thomas Scientific #1141J52) or detected using the LI-COR Odyssey Fc system. PrP^C^ blots were reprobed with the rabbit polyclonal anti-actin antibody 20-33 (1:10,000; Millipore Sigma #A5060).

### Prion strains

Hamster prion strains 263K, 139H, Hyper (HY), and Drowsy (DY) were originally obtained as 10% (w/v) brain homogenates in PBS from Jason Bartz (Creighton University). To generate a renewable source of the 263K, HY, and 139H strains, CAD5-PrP^-/-^ cells expressing HaPrP were used (42). The mouse prion strains RML and 22L were obtained from terminally ill non-transgenic C57BL/6 mice. Brains of prion-infected mice were homogenized in PBS to generate a 10% (w/v) brain homogenate stock, which was aliquoted and stored at -80 °C. The homogenization step was conducted using a Minilys homogenizer (Bertin Technologies) with CK14 soft tissue homogenization tubes containing 1.4 mm zirconium oxide beads (Bertin Technologies #P000912-LYSK0-A). Brains were homogenized using 3 cycles of 60 s at maximum speed with each cycle followed by 10 min of incubation on ice. For cellular homogenates, prion-infected CAD5-PrP^-/-^ cells expressing HaPrP were cultured in 10-cm tissue culture plates. After reaching confluency, the cells were washed in cold PBS and then scraped into a small volume of PBS and collected into CK14 soft tissue homogenization tubes supplemented with additional 0.5 mm zirconia beads (BioSpec #11079105Z). The cells were homogenized using 3 cycles of 60 s at maximum speed, with a 10 min incubation on ice between each cycle. Total protein concentrations in cell lysates were quantified using the BCA assay.

### Cellular prion infections

Stably transfected CAD5-PrP^-/-^ cells were plated in either 12- or 24-well dishes at a density of 150,000 or 100,000 cells/well respectively. The following day, cells were challenged with 0.1% (w/v) prion-infected brain homogenate or 100 µg of prion-infected cellular homogenate. After 48-72 h of incubation (depending on cell confluency), the cells were washed 2X with PBS and then passaged at 1:3 dilution into 12-well plates. The cells were then passaged six times in 12-well plates to allow for *de novo* prion accumulation. At the sixth passage, the cells were scaled up to 6-cm dishes for lysis and analysis of successful prion infection.

### Proteinase K digestions

To determine the infection status of cell lines challenged with prions, cells were lysed using lysis buffer, as described above, and then lysates were digested with proteinase K (PK). After determining protein concentration using the BCA assay, 0.5 to 1 mg of cell lysate was digested with PK (ThermoFisher Scientific #EO0491) at a concentration of 50 µg/mL and a PK:protein ratio of 1:50. The digestion was carried out at 37 °C for 1 h with shaking at 600 rpm. The digestion was halted by the addition of PMSF to a final concentration of 2 mM, and then sarkosyl (Millipore Sigma #61747) was added to a final concentration of 2% (v/v). Digested samples were ultracentrifuged at 100,000x *g* for 1 h at 4 °C using a TLA-55 rotor (Beckman Coulter) to concentrate the insoluble, PK-resistant fraction. The supernatant was discarded, and the pellet was resuspended in 25 µL of 1X Bolt LDS sample buffer, boiled at 95 °C, and then stored at -80 °C for subsequent analysis by immunoblotting.

### Purification of recombinant PrP

Recombinant untagged full-length PrPs (residues 23-230 for mouse PrP, 23-231 for hamster PrP and BVPrP, residues 25-233 for elk PrP and ovine PrP) were expressed and purified as previously described (42, 44, 49). Briefly, sequences were inserted into the pET-41 expression vector and then proteins were expressed in *E.coli* Rosetta2(DE3) cells using an autoinduction system. For BVPrP, the M109 variant was utilized and for sheep PrP, the ARQ allele was used. Cells were lysed using BugBuster Master Mix solution (Millipore Sigma #71456) and then inclusion bodies were isolated. Inclusion bodies were solubilized in 8 M guanidine hydrochloride (GdnHCl) and then PrP was captured using Ni-NTA Superflow beads (Qiagen #30410). Bead-bound PrP was refolded on-column using a 4 h gradient from 6 to 0 M GdnHCl and then eluted using a gradient from 0 to 500 mM imidazole. PrP-containing fractions were dialyzed into 10 mM sodium phosphate pH 5.8, filtered through a 0.22-micron filter, and then stored at -80 °C. The concentration of purified recombinant PrP was determined through absorbance at 280 nm using a NanoDrop spectrophotometer. The purity of recombinant PrP was assessed through SDS-PAGE followed by staining with Coomassie brilliant blue. To confirm the successful refolding of recombinant PrP to an α-helical state, circular dichroism experiments on protein dialyzed in 10 mM sodium phosphate pH 7.3 buffer were performed at 22 °C using a JASCO J-715 spectropolarimeter.

### Thioflavin T aggregation assays

Recombinant PrP was dialyzed in 10 mM sodium phosphate pH 7.3 buffer overnight or for 4 h at 4 °C with two buffer changes. To remove any pre-existing aggregates, the dialyzed protein was ultracentrifuged (100,000x *g*) for 1 h at 4 °C. The reaction mixtures were prepared as follows: 0.1 mg/mL recombinant PrP, 10 µM Thioflavin T (ThT), and 135 mM NaCl in a total volume of 100 µL of 10 mM sodium phosphate pH 7.3. The mixtures were plated in black, clear-bottom 96-well plates (ThermoFisher Scientific #265301) and then sealed with a clear film (VWR #89134-428). The plate was then placed in a BMG CLARIOstar plate reader and subjected to cycles of 4 min shaking (double orbital, 700 rpm) and 1 min rest/read at 37 °C. The ThT readings (excitation 444 ± 5 nm; emission 485 ± 5 nm) were taken every 5 min during the rest period using a gain setting of 1600. For the experiments involving the C-terminal PrP chimeras, the following settings were used to prevent the ThT signal from approaching the detection limit of the instrument: excitation 444 ± 4 nm; emission 485 ± 4 nm; gain = 1500. The lag phases for the aggregation reactions were quantified as previously described (44). To determine the maximum:plateau fluorescence ratio for each replicate, the maximum ThT fluorescence achieved during the experimental timeframe was divided by the ThT fluorescence at the experimental endpoint (83.25 h). Samples that did not aggregate within the experimental timeframe were excluded from the lag phase, maximum ThT fluorescence, and maximum:plateau ThT fluorescence comparisons.

### Statistical analysis

All statistical analysis was performed using GraphPad Prism (version 10.1.1) with a statistical threshold of P < 0.05. Lag phases, maximum ThT fluorescence values, and maximum:plateau ThT fluorescence ratios were compared using one-way ANOVA followed by either Dunnett’s (when comparing all samples to recombinant wild-type BVPrP) or Tukey’s (when performing all possible pairwise comparisons) multiple comparisons tests.

## Results

### The role of the BVPrP N-terminal domain and N-glycosylation in cross-species prion infection

We have previously demonstrated that monoclonal CAD5-PrP^-/-^ cell lines stably expressing BVPrP are susceptible to infection with several different strains of mouse and hamster prions (44). Indeed, following three passages post-infection, a monoclonal line of CAD5-PrP^-/-^(BVPrP) cells exhibits robust levels of PK-resistant PrP (PrP^res^), indicative of PrP^Sc^ production (**Fig. S1**). Monoclonal lines are ideal for cellular prion infection experiments as they obviate the need for including G418 in the culture medium, which helps to maintain PrP^C^ expression levels but also interferes with *de novo* prion infection (49). However, polyclonal cell lines stably expressing BVPrP and cultured in the presence of a low concentration of G418 (0.2 mg/mL) can also become infected following challenge with mouse and hamster prion strains, although they display comparatively lower levels of PrP^res^ at passage three (**Fig. S1**). To facilitate studying the susceptibility of many PrP variants to cross-species prion infection, we chose to use polyclonal lines of stably transfected CAD5-PrP^-/-^ cells for our experiments and analyzed them following six passages post-infection to allow sufficient time for PrP^res^ accumulation. Although both the M109 and I109 variants of BVPrP are susceptible to cross-species prion infection when expressed in cultured cells (44), we chose to exclusively use the M109 variant for this study.

While not essential for prion replication to occur, post-translational modification by N-glycans and the disordered N-terminal domain can modulate the conversion of PrP^C^ into PrP^Sc^ and the resultant prion strain properties (50–55). To test the roles of the N-terminal region and N-glycosylation in the ability of BVPrP to enable cross-species prion replication, we generated BVPrP mutants lacking portions of the N-terminal domain as well as mutants with asparagine to glutamine substitutions that abolish the N-glycan attachment sites (**Fig. S2A**). All mutants could be successfully expressed in CAD5-PrP^-/-^ cells (**Fig. S2B**). The mutant BVPrP-expressing cell lines were tested for their ability to replicate two distinct strains of mouse prions (RML and 22L) and two distinct strains of hamster prions (263K and HY), in comparison to CAD5-PrP^-/-^ cells expressing wild-type BVPrP. Mutants lacking the complete N-terminal domain (Δ23-89) as well a mutant lacking the second N-glycan attachment site (N197Q) formed PrP^res^ following challenge with either mouse or hamster prions (**Fig. S2C-F**). In contrast, cells expressing mutant BVPrP lacking the first N-glycan site (N181Q), either alone or in combination with the N197Q mutation, did not accumulate PrP^res^ following prion infection. These results indicate that the N-terminal domain of BVPrP as well as N-glycosylation at residue N197 are dispensable for cross-species prion replication.

### Identification of key residues in BVPrP that enable cross-species prion replication using chimeric mouse/bank vole PrPs

The sequence of mature MoPrP lacking the N- and C-terminal signal sequences differs from the sequence of BVPrP at eight positions (**Fig. 1A**). These amino acid residues are present throughout the sequence of BVPrP, with three residues found in the disordered N-terminal domain (G54, G72, G80), three residues in the central globular domain (M109, N155, N170), and two residues near the C-terminal GPI anchor attachment site (E227 and S230). To investigate which of these residues are critical for enabling cross-species prion replication, three Mo/BVPrP chimeras were initially generated. Mouse chimera 1 (MoC1) consists of MoPrP with the three BVPrP-specific residues from the N-terminal domain, MoC2 contains the three BVPrP residues from the central globular domain, and MoC3 contains the two BVPrP residues near the GPI anchor attachment site (**Fig. 1A**). As CAD5-PrP^-/-^ cells expressing wild-type MoPrP are resistant to hamster prions (42, 44), our goal was to identify combinations of BVPrP residues that enable replication of hamster prions without losing the ability to become infected by mouse prions. The three chimeras were stably expressed in CAD5-PrP^-/-^ cells and exhibited similar levels of PrP expression to wild-type BVPrP (**Fig. 1B**). Cells expressing the chimeras were challenged with three different hamster prion strains: 263K, HY, and 139H. Cells expressing wild-type MoPrP or BVPrP were used as negative and positive controls, respectively. Of the three chimeras, only cells expressing MoC2 were able to be infected by the three hamster prion strains, as indicated by the presence of PrP^res^ in cell lysates (**Fig. 1C-E**). Thus, residues M109, N155, and N170 of BVPrP enable cross-species prion replication when inserted into the sequence of MoPrP.

**Figure 1.**
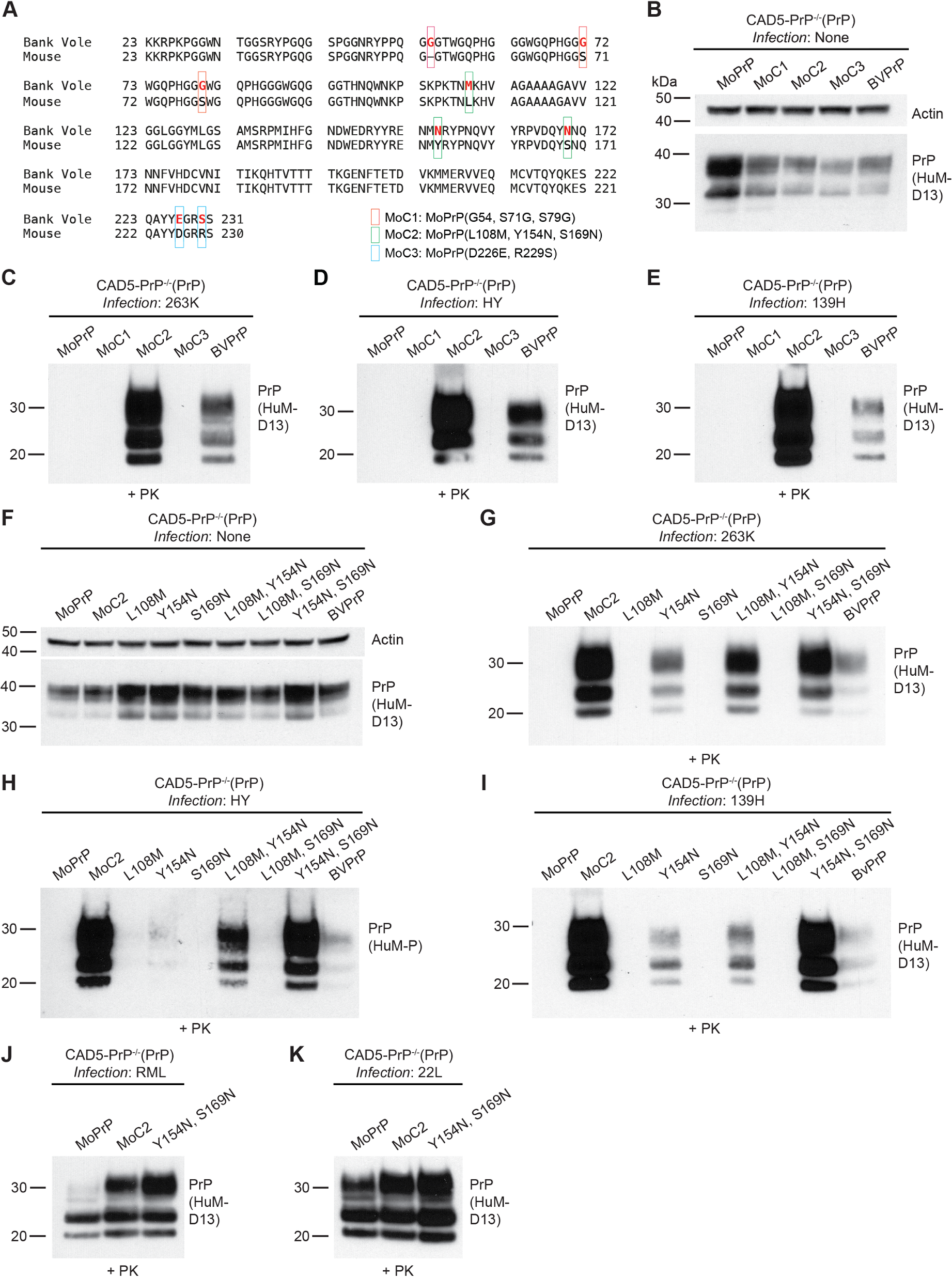
Identification of BVPrP residues that render MoPrP susceptible to hamster prions. **A**) Amino acid sequence alignment of the sequences of BVPrP and MoPrP following removal of N- and C-terminal signal sequences. The sequences differ at eight positions (depicted by red lettering in the BVPrP sequence). The location of residue substitutions in the MoC1, MoC2, and MoC3 chimeras is indicated. **B**) Immunoblot for PrP^C^ in undigested lysates from CAD5-PrP^-/-^ cells stably expressing the indicated wild-type or chimeric PrP molecules. The blot was reprobed with an antibody against actin. **C-E**) Immunoblots of PrP^res^ levels in lysates from CAD5-PrP^-/-^ cells stably expressing the indicated PrP molecules challenged with either 263K (**C**), HY (**D**), or 139H (**E**) hamster prions. **F**) Immunoblot for PrP^C^ in undigested lysates from CAD5-PrP^-/-^ cells stably expressing the indicated wild-type or mutant PrP molecules. The blot was reprobed with an antibody against actin. **G-I**) Immunoblots of PrP^res^ levels in lysates from CAD5-PrP^-/-^ cells stably expressing the indicated PrP molecules challenged with either 263K (**G**), HY (**H**), or 139H (**I**) hamster prions. **J-K**) Immunoblots of PrP^res^ levels in lysates from CAD5-PrP^-/-^ cells stably expressing wild-type MoPrP, the MoC2 chimera, or Y154N/S169N-mutant MoPrP challenged with RML (**J**) or 22L (**K**) mouse prions. PrP^C^ and PrP^res^ were detected using the antibodies HuM-D13 or HuM-P. In all panels, the molecular weight markers indicate kDa.

To determine which residues within MoC2 were sufficient for permitting cross-species prion replication, each possible combination of the three residues was tested. The resultant six new cell lines exhibited roughly similar levels of PrP expression (**Fig. 1F**). These cell lines were challenged with the same three hamster prion strains as above. We found that the addition of BVPrP residues N155 and N170 (Y154N and S169N mutations in MoPrP) produced levels of PrP^res^ similar to those observed in cells expressing MoC2 for all three hamster prion strains tested (**Fig. 1G-I**). Of the two residues, N155 appears to be more important as it permitted infection with 263K and 139H prions in isolation, albeit with lower levels of PrP^res^ produced, whereas cells expressing a mutant MoPrP containing just the N170 substitution were resistant to all three hamster prion strains. Importantly, cells expressing either the MoC2 or N155/N170 mutants retained their ability to be infected with two different strains of mouse prions: RML and 22L (**Fig 1J-K**). Therefore, the addition of BVPrP residues N155 and N170 enable MoPrP to function as a permissive substrate for cross-species prion replication, akin to wild-type BVPrP.

To further investigate the importance of these two residues, a mutant was generated in which the corresponding mouse residues were substituted into the sequence of BVPrP (N155Y and N170S mutations). In CAD5-PrP^-/-^ cells, N155Y/N170S-mutant BVPrP expressed at similar levels to wild-type BVPrP (**Fig. 2A**). When N155Y/N170S-mutant cells were infected with mouse (RML, 22L) or hamster (263K, HY, DY) prions, little to no PrP^res^ was observed, suggesting that the cells were resistant to prion infection (**Fig. 2B, C**). In contrast, abundant PrP^res^ was observed in cells expressing wild-type BVPrP following challenge with each of the four prion strains. To further test the resistance of this BVPrP mutant to prion infection, RML and 263K prions adapted to the BVPrP sequence by passaging through cells expressing wild-type BVPrP (BV.RML and BV.263K prions) were used as the prion inoculum. Despite the use of BVPrP-adapted prion strains, cells expressing N155Y/N170S-mutant BVPrP were still resistant to prion infection (**Fig. 2D**). Thus, asparagine residues 155 and 170 of BVPrP are necessary for cross-species prion replication. Consistent with our findings, the importance of these asparagine residues for the unusual properties of BVPrP has previously been inferred from prion infection studies in phylogenetically related rodent species as well as from *in vitro* prion conversion experiments (17, 35).

**Figure 2.**
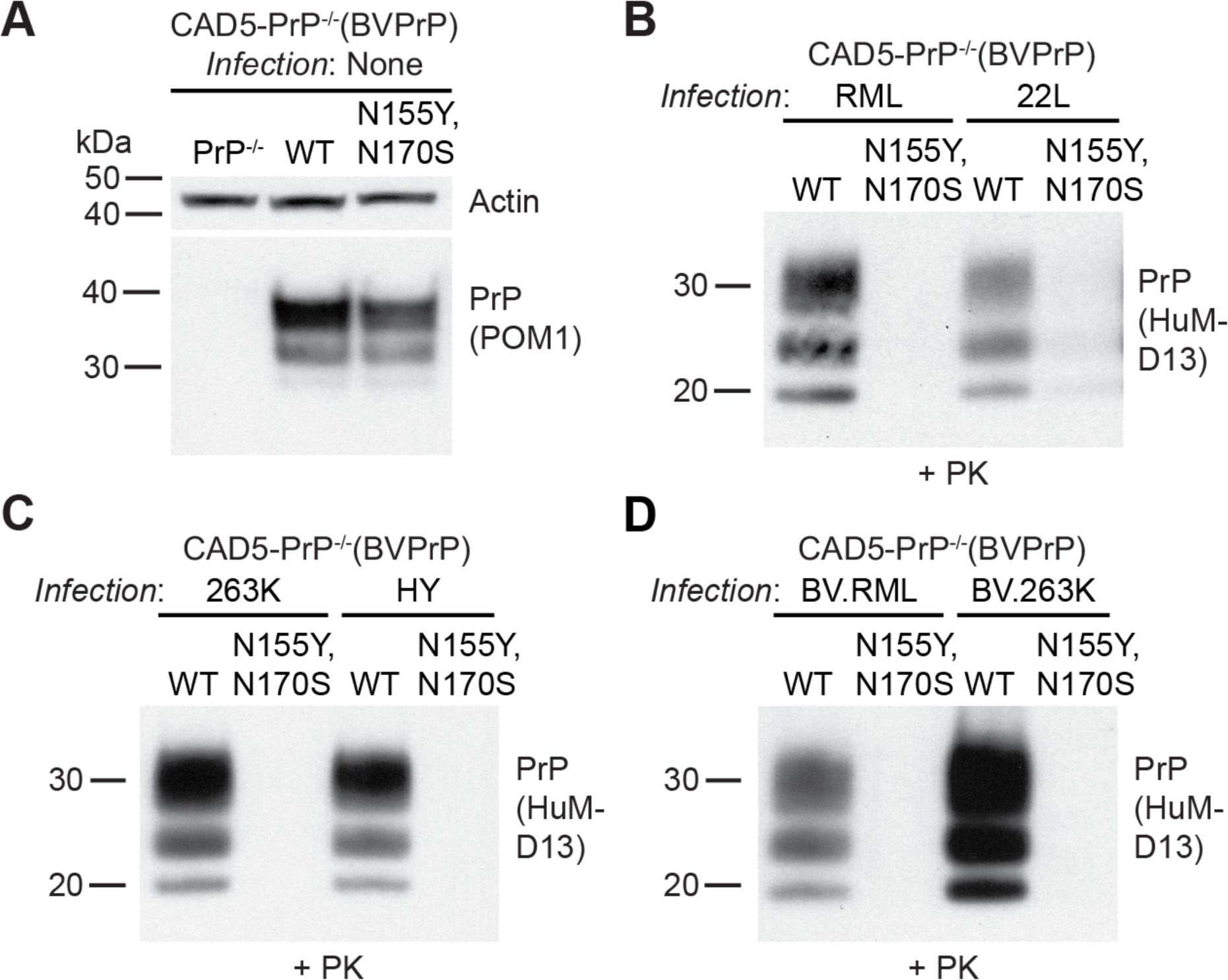
Asparagine residues at positions 155 and 170 of BVPrP are necessary for cross-species prion replication. **A**) Immunoblot for PrP^C^ in undigested lysates from CAD5-PrP^-/-^ cells stably expressing the indicated BVPrP molecules. Lysate from untransfected CAD5-PrP^-/-^ cells is included as a negative control. The blot was reprobed with an antibody against actin. **B-D**) Immunoblots of PrP^res^ levels in lysates from CAD5-PrP^-/-^ cells stably expressing either wild-type or N155Y/N170Q-mutant BVPrP challenged with mouse (**B**), hamster (**C**), or bank vole-adapted (**D**) prion strains. PrP^C^ was detected using the antibody POM1 whereas PrP^res^ was detected using HuM-D13. In all panels, the molecular weight markers indicate kDa.

### Identification of additional residues in BVPrP that enable cross-species prion replication using chimeric hamster/bank vole PrPs

Our results demonstrate that asparagine residues at positions 155 and 170 of BVPrP are critical for cross-species prion replication. However, the sequence of HaPrP also contains these two residues, yet HaPrP is unable to enable cross-species prion replication like BVPrP (44). Thus, BVPrP must contain additional residues that are absent in HaPrP that allow it to act as a permissive substrate for cross-species prion replication. The mature sequence of HaPrP differs from BVPrP at only seven residues (**Fig. 3A**). To investigate which BVPrP residues are critical, we generated two Ha/BVPrP chimeras. Hamster chimera 1 (HaC1) consists of HaPrP with five BVPrP-specific residues from the central globular domain whereas HaC2 consists of HaPrP with the two BVPrP residues near the GPI anchor attachment site (**Fig. 3A**). The two chimeras, along with HaPrP as a negative control and BVPrP as a positive control, were stably expressed in CAD5-PrP^-/-^ cells (**Fig. 3B**). To determine which combination of BVPrP residues enable HaPrP to replicate mouse prions, cells were treated with the RML and 22L strains. Cells expressing either HaC1 or wild-type BVPrP could be successfully infected with both prion strains, with HaC1-expressing cells displaying higher levels of PrP^res^ than cells expressing BVPrP (**Fig. 3C, D**).

**Figure 3.**
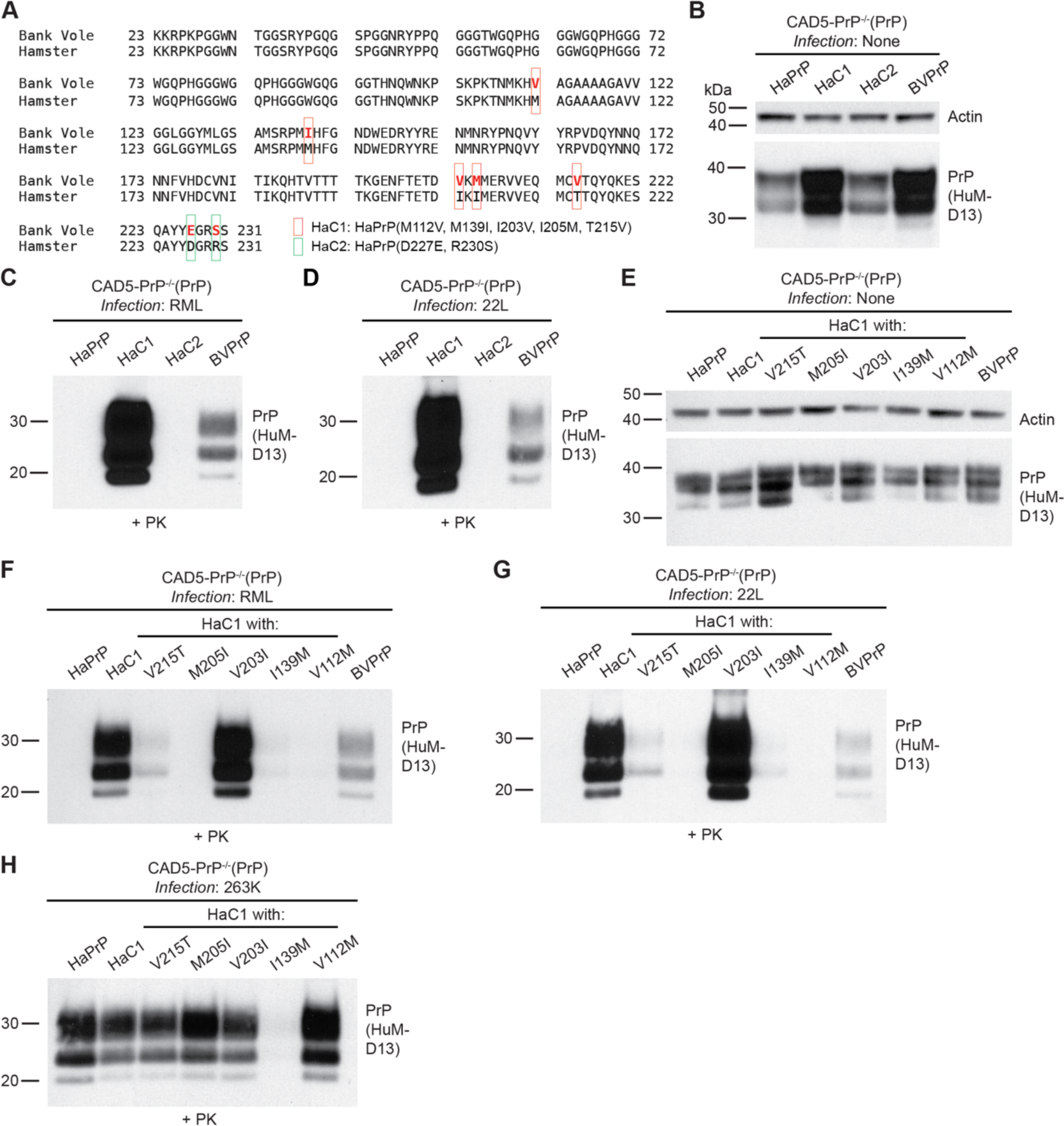
Identification of BVPrP residues that render HaPrP susceptible to mouse prions. **A**) Amino acid sequence alignment of the sequences of BVPrP and HaPrP following removal of N- and C-terminal signal sequences. The sequences differ at seven positions (depicted by red lettering in the BVPrP sequence). The location of residue substitutions in the HaC1 and HaC2 chimeras is indicated. **B**) Immunoblot for PrP^C^ in undigested lysates from CAD5-PrP^-/-^ cells stably expressing the indicated wild-type or chimeric PrP molecules. The blot was reprobed with an antibody against actin. **C-D**) Immunoblots of PrP^res^ levels in lysates from CAD5-PrP^-/-^ cells stably expressing the indicated PrP molecules challenged with either RML (**C**) or 22L (**D**) mouse prions. **E**) Immunoblot for PrP^C^ in undigested lysates from CAD5-PrP^-/-^ cells stably expressing the indicated wild-type or mutant PrP molecules. The blot was reprobed with an antibody against actin. **F-H**) Immunoblots of PrP^res^ levels in lysates from CAD5-PrP^-/-^ cells stably expressing the indicated PrP molecules challenged with either RML (**F**), 22L (**G**), or 263K (**H**) prions. PrP^C^ and PrP^res^ were detected using the antibody HuM-D13. In all panels, the molecular weight markers indicate kDa.

To determine which of the BVPrP residues in HaC1 were critical for enabling cross-species prion replication, each individual BVPrP residue substitution was generated independently in HaPrP and stably expressed in CAD5-PrP^-/-^ cells (**Fig. S3A**). The cell lines were then challenged with RML and 22L prions. Only cells expressing wild-type BVPrP or the HaC1 chimera showed any signs of infection, indicating that none of five BVPrP residues are individually sufficient to enable HaPrP to function as a permissive substrate for cross-species prion replication (**Fig. S3B, C**). Therefore, an alternate approach was taken. We hypothesized that individual residue reversions within the HaC1 chimera to the original HaPrP residue would reveal the importance of each of the five distinct residues. Lines of stably transfected CAD5-PrP^-/-^ cells were generated, with each PrP variant expressing at roughly similar levels (**Fig. 3E**). The lines were then challenged with RML and 22L prions. The V203I substitution did not affect the ability of HaC1 to become infected with mouse prions, indicating that BVPrP residue 203 is dispensable for cross-species prion infection (**Fig. 3F, G**). Importantly, cells expressing either HaC1 or HaC1 with the V203I substitution retained their susceptibility to hamster 263K prions, indicating that the identified BVPrP residues enable cross-species prion replication and that observed effects are not simply due to sequence similarity between PrP^C^ and PrP^Sc^ (**Fig. 3H**). Lower amounts of PrP^res^ were also observed following infection of cells expressing HaC1 with the V215T or I139M substitutions with mouse prions (**Fig. 3F, G**). Cells expressing HaC1 with the V215T substitution, but not the I139M substitution, were also susceptible to hamster prions, indicating that, like residue 203, residue 215 of BVPrP is not critical for cross-species prion replication (**Fig. 3H**).

A Ha/BVPrP chimera consisting of HaPrP with BVPrP-specific residues at positions 112, 139, 205, and 215, termed HaC3, was used as a starting point to more precisely map the BVPrP residues that enable HaPrP to serve as a substrate for cross-species prion replication (**Fig. 4A**). All possible two- and three-residue combinations of these four positions were generated and stably expressed in CAD5-PrP^-/-^ cells (**Fig. 4B, C**). When challenged with RML prions, only the HaPrP chimera containing the M112V, M139I, and I205M substitutions became infected with prions (**Fig. 4D**). The same chimera was also susceptible to both mouse 22L and hamster 263K prions (**Fig. 4E, F**). While no two-residue combination was sufficient to enable replication of RML prions, the addition of BVPrP residues V112 and I139 to HaPrP did permit robust replication of the 22L strain (**Fig. 4E**). Thus, we conclude that three BVPrP residues (V112, I139, and M205) are sufficient for permitting HaPrP to replicate both mouse and hamster prions, although certain strains may require only a subset of these residues.

**Figure 4.**
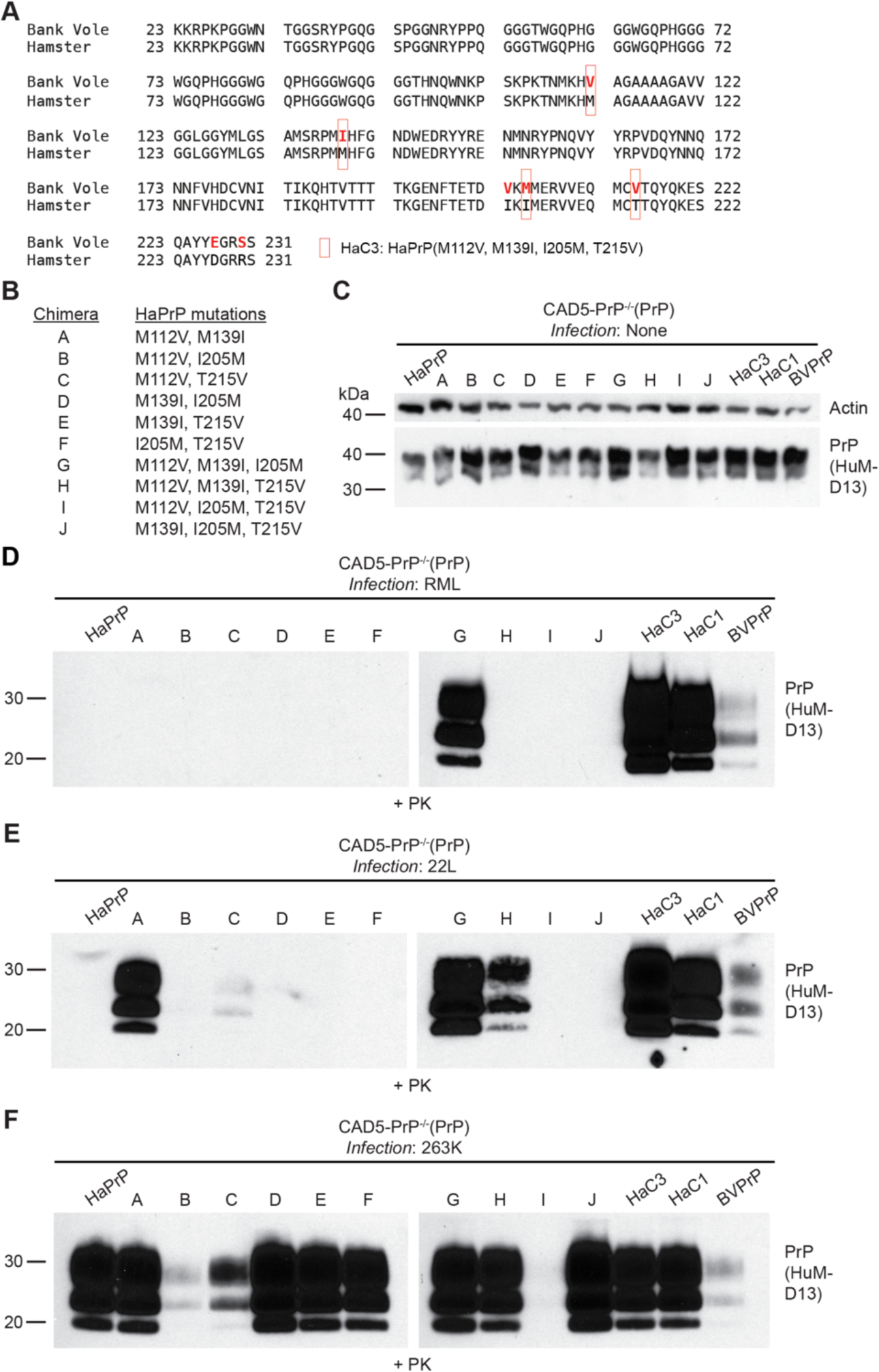
Addition of three key BVPrP residues to HaPrP permits cross-species prion replication. **A**) Amino acid sequence alignment of the sequences of BVPrP and HaPrP with the location of residue substitutions in the HaC3 chimera indicated. **B**) Chimeras A-J contain various combinations of BVPrP residues 112, 139, 205, and 215 inserted into the sequence of HaPrP. **C**) Immunoblot for PrP^C^ in undigested lysates from CAD5-PrP^-/-^ cells stably expressing the indicated wild-type or chimeric PrP molecules. The blot was reprobed with an antibody against actin. **D-F**) Immunoblots of PrP^res^ levels in lysates from CAD5-PrP^-/-^ cells stably expressing the indicated PrP molecules challenged with either RML (**D**), 22L (**E**), or 263K (**F**) prions. For each strain, all samples were transferred to the same membrane. PrP^C^ and PrP^res^ were detected using the antibody HuM-D13. In all panels, the molecular weight markers indicate kDa.

### Identification of a C-terminal motif in BVPrP that impedes cross-species prion replication

BVPrP only contains two unique residues that are not found in the sequences of either MoPrP or HaPrP: residues E227 and S230 located near the GPI anchor attachment site. These two residues have been implicated in BVPrP’s propensity to misfold spontaneously and its ability to function as a permissive substrate for prion replication (36, 37). Unexpectedly, we found that chimeric PrPs that lack BVPrP residues E227 and S230 function as better substrates for cross-species prion replication than wild-type BVPrP, based on the higher levels of PrP^res^ observed following infection with hamster or mouse prions (**Fig. 1C-E, G-I; Fig. 3C, D, F, G**; **Fig. 4D-F**). To further investigate the role of these two residues in cross-species prion replication, we re-tested the chimeric mouse and hamster PrPs containing BVPrP residues E227 and S230, previously referred to as MoC3 and HaC2. New lines of stably transfected CAD5-PrP^-/-^ cells were generated, along with cells expressing wild-type MoPrP or HaPrP as controls (**Fig. 5A, B**).

**Figure 5.**
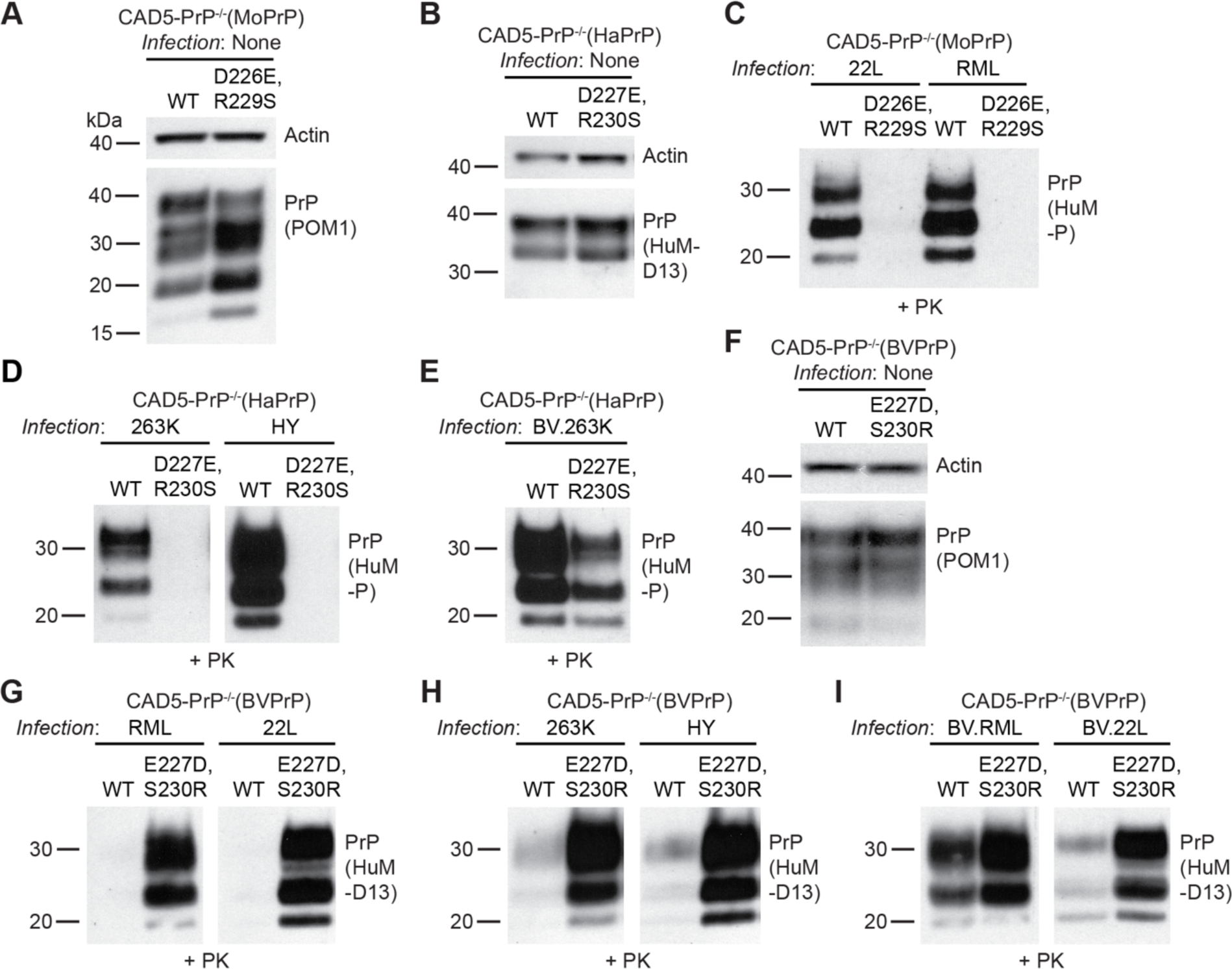
Residues E227 and S230 in BVPrP impede cross-species prion infection in cultured cells. **A**) Immunoblot for PrP^C^ in undigested lysates from CAD5-PrP^-/-^ cells stably expressing wild-type or D226E/R229S-mutant MoPrP. **B**) Immunoblot for PrP^C^ in undigested lysates from CAD5-PrP^-/-^ cells stably expressing wild-type or D227E/R230S-mutant HaPrP. **C**) Immunoblot of PrP^res^ levels in lysates from CAD5-PrP^-/-^ cells stably expressing either wild-type or D226E/R229S-mutant MoPrP challenged with 22L or RML mouse prions. **D-E**) Immunoblots of PrP^res^ levels in lysates from CAD5-PrP^-/-^ cells stably expressing either wild-type or D227E/R230S-mutant HaPrP challenged with 263K, HY (**D**), or BV.263K (**E**) prions. **F**) Immunoblot for PrP^C^ in undigested lysates from CAD5-PrP^-/-^ cells stably expressing wild-type or E227D/S230R-mutant BVPrP. **G-I**) Immunoblots of PrP^res^ levels in lysates from CAD5-PrP^-/-^ cells stably expressing either wild-type or E227D/S230R-mutant BVPrP challenged with mouse (**G**), hamster (**H**), or bank vole-adapted (**I**) prion strains. PrP^C^ was detected using the antibodies POM1 or HuM-D13 whereas PrP^res^ was detected using HuM-P or HuM-D13. The blots in panels A, C, and F were reprobed with an antibody against actin. In all panels, the molecular weight markers indicate kDa.

Cells expressing MoPrP containing the D226E and R229S substitutions were resistant to RML and 22L prions, indicated by a lack of PrP^res^ following prion infection, whereas cells expressing wild-type MoPrP were readily infected (**Fig. 5C**). Similarly, cells expressing HaPrP containing the D227E and R230S substitutions were resistant to 263K and HY hamster prions (**Fig. 5D**). CAD5-PrP^-/-^ cells expressing HaPrP containing the D227E and R230S substitutions accumulated less PrP^res^ following infection with BV.263K prions (**Fig. 5E**), suggesting that decreased sequence identity between the C-terminal chimeric PrP^C^ substrates and PrP^Sc^ in the inoculum is not sufficient to explain the observed effects, as BV.263K prions are more closely sequence matched to the D227E/R230S mutant than wild-type HaPrP. We could not test whether the same is true when using BV.RML prions, since cells expressing wild-type MoPrP are resistant to infection by BVPrP-adapted mouse prion strains (44).

We also conducted the opposite experiment in which the two C-terminal residues in BVPrP were substituted with the equivalent residues that are common to the sequences of both MoPrP and HaPrP. A BVPrP variant with the E227D and S230R substitutions was generated and stably expressed in CAD5-PrP^-/-^ cells (**Fig. 5F**). When these cells were challenged with mouse and hamster prion strains, they exhibited much higher levels of PrP^res^ than cells expressing wild-type BVPrP (**Fig. 5G, H**). Once again, sequence similarity between PrP^C^ and PrP^Sc^ was not the driving force for this observation since higher levels of PrP^res^ were observed in cells expressing E227D/S230R-mutant BVPrP following infection with BVPrP-adapted RML and 22L prions compared to wild-type BVPrP-expressing cells (**Fig. 5I**). Therefore, the C-terminal region of BVPrP contains amino residues that interfere with prion replication.

To further assess the inhibitory properties associated with BVPrP residues E227 and S230 and how they relate to sequence similarity, we compared four BVPrP substrates harboring various amino acids at positions 227 and 230. These include BVPrP with the wild-type residues (E,S), mouse/hamster equivalent residues (D,R), human PrP residues (Q,S) or residues found in the sequence of elk/ovine/bovine/rabbit/dog PrP (Q,A) (**Fig. 6A**). The BVPrP variants were stably expressed in CAD5-PrP^-/-^ cells, with roughly equal levels of PrP^C^ expression between the four lines (**Fig. 6B**). When these cells were challenged with mouse (RML, 22L) or hamster (263K, HY, DY) prions, cells expressing wild-type BVPrP accumulated the lowest amounts of PrP^res^ (**Fig. 6C, D**). Simply substituting BVPrP residue E227 with the Q227 residue found in human PrP drastically improved the ability of BVPrP to permit cross-species prion replication. Notably, Q227 is not present in any of the mouse or hamster PrP^Sc^ inocula, confirming that the differences in PrP^res^ accumulation are not mediated by sequence similarity between PrP^C^ and PrP^Sc^. Moreover, identical results were obtained with four different strains of BVPrP-adapted prions (**Fig. 6E**). In summary, the presence of residues E227 and S230 in wild-type BVPrP substantially reduces susceptibility to cross-species prion infection and the effect is independent of sequence similarity.

**Figure 6.**
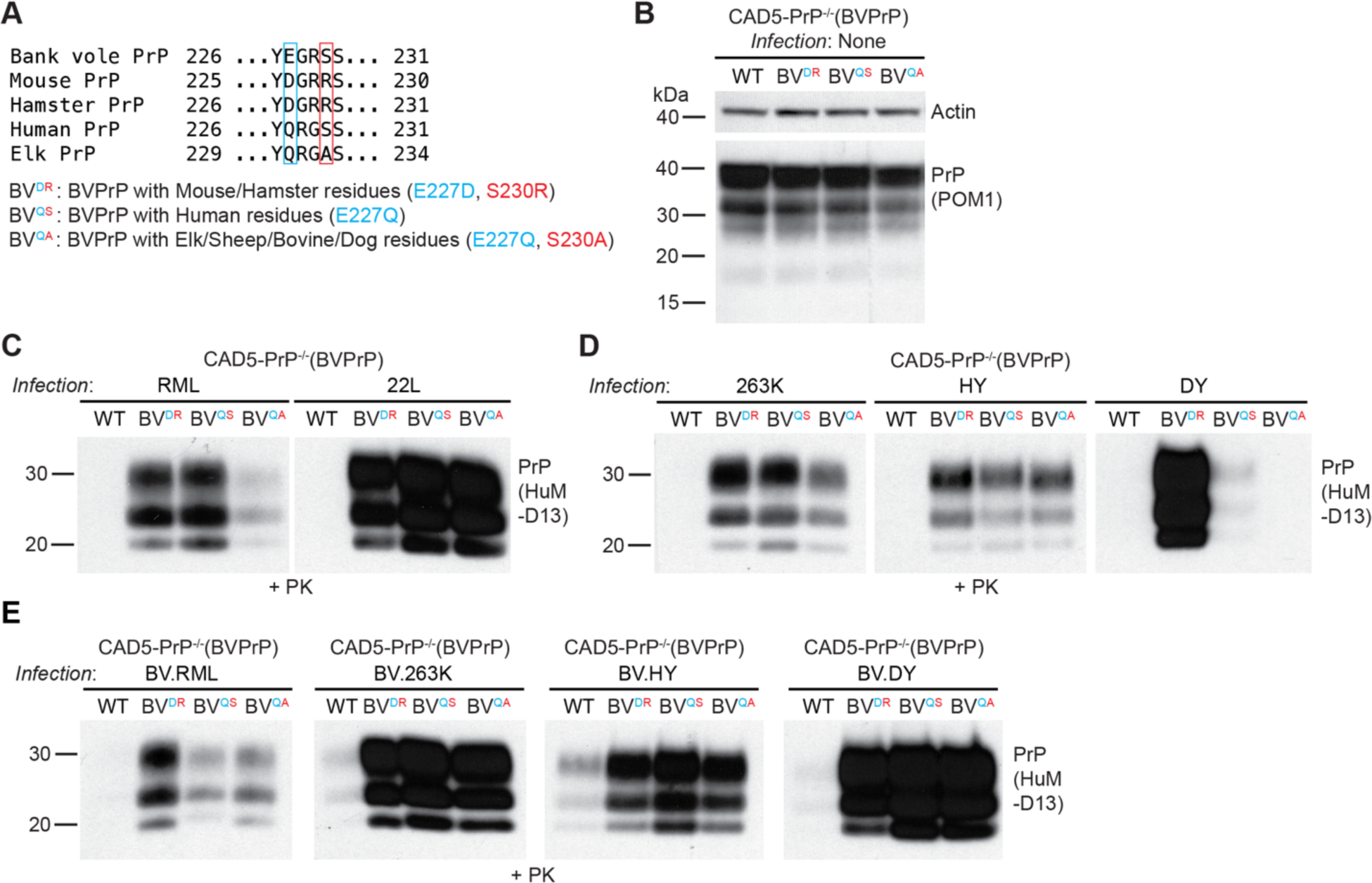
A C-terminal motif in BVPrP interferes with cross-species prion replication independent of sequence similarity. **A**) Amino acid sequence alignment of the sequences of the C-terminal regions of bank vole, mouse, hamster, human, and elk PrP. A description of the C-terminal BVPrP chimeras is also given. **B**) Immunoblot for PrP^C^ in undigested lysates from CAD5-PrP^-/-^ cells stably expressing the indicated wild-type or chimeric BVPrP molecules. The blot was reprobed with an antibody against actin. **C-E**) Immunoblots of PrP^res^ levels in lysates from CAD5-PrP^-/-^ cells stably expressing the indicated PrP molecules challenged with either mouse (**C**), hamster (**D**), or bank vole-adapted (**E**) prion strains. PrP^C^ was detected using the antibody POM1 whereas PrP^res^ was detected using HuM-D13. In all panels, the molecular weight markers indicate kDa.

### PrP aggregation kinetics mimic the relative permissiveness of different PrPs to cross-species prion replication

To investigate the mechanism by which BVPrP may permit cross-species prion replication, we generated and purified recombinant (r) full-length untagged PrPs from several different species including bank vole, mouse, hamster, sheep, and elk (**Fig. 7A**). Circular dichroism spectroscopy confirmed that each rPrP was successfully refolded to an α-helical state, consistent with the structure of PrP^C^ (**Fig. 7B**). We then compared the relative propensities of the rPrPs to spontaneously polymerize into aggregates by performing a ThT aggregation assay at 37 °C with continuous shaking. The ThT assays were performed in a physiological buffer containing 135 mM salt and a pH of 7.3. rBVPrP formed spontaneous aggregates much more rapidly than the rPrPs from the other four species (**Fig. 7C**). Indeed, the lag phase for rBVPrP aggregation was significantly shorter than the lag phases for aggregation of hamster, ovine, and elk PrP (**Fig. 7D**). rMoPrP displayed intermediate aggregation kinetics, but consistently polymerized more slowly than rBVPrP (**Fig. 7C, D**). We also found that the aggregates formed by rBVPrP exhibited higher ThT fluorescence than those composed of the other rPrPs (**Fig. 7C, E**). The shapes of the aggregation curves also differed, with rBVPrP initially reaching a higher maximum ThT fluorescence before plateauing at a lower value, whereas the other rPrPs plateaued at ThT fluorescence values similar to the maximum ThT signal attained (**Fig. 7C, F**). The underlying explanation for the differences in curve shape and plateau fluorescence remains to be determined but could be related to structural differences among the aggregates and/or distinct aggregation pathways. These results demonstrate that the ability to permit cross-species prion replication is correlated with enhanced spontaneous aggregation kinetics of rPrP.

**Figure 7.**
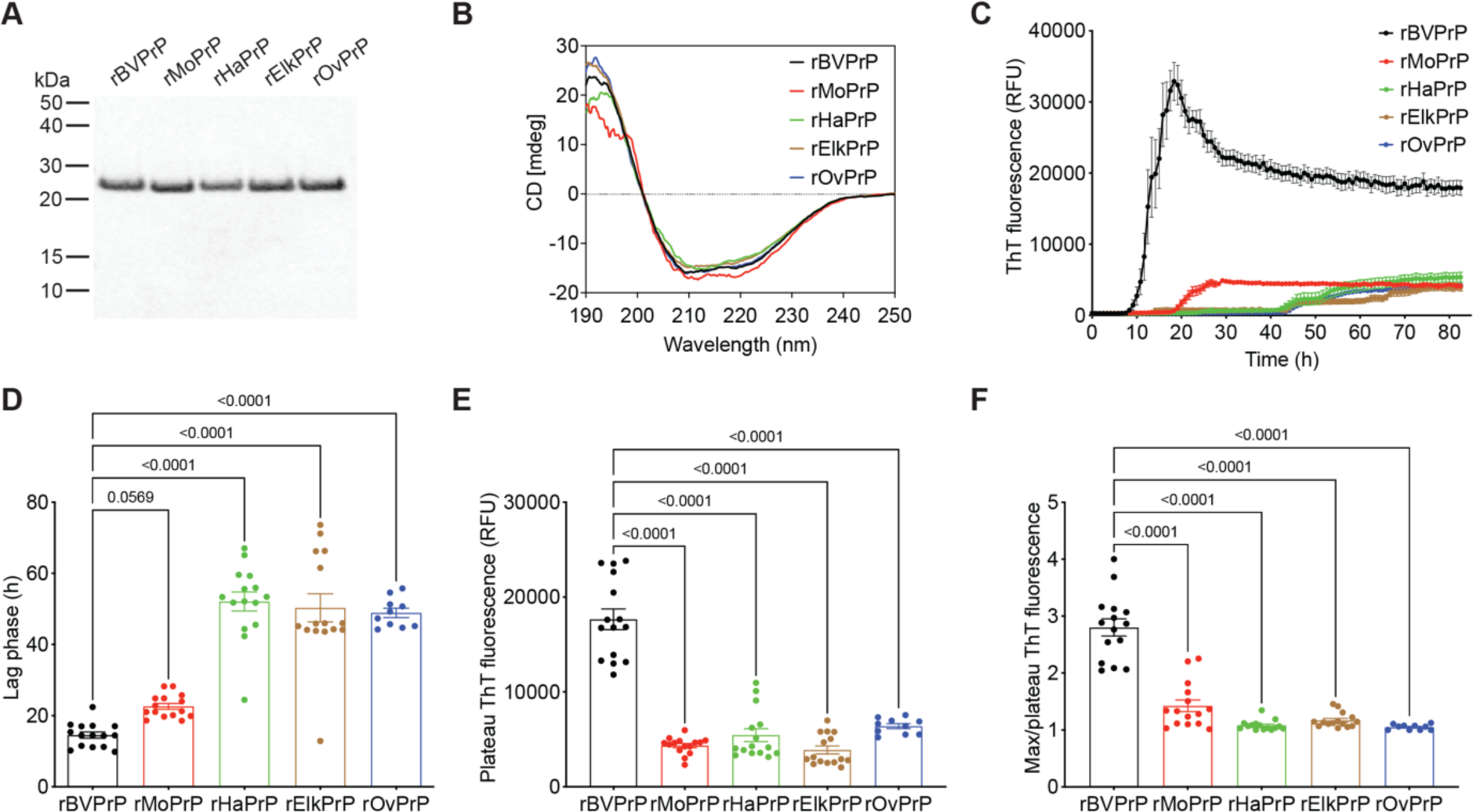
Spontaneous aggregation kinetics of recombinant PrPs from different species. A) SDS-PAGE followed by Coomassie blue staining of purified recombinant BV, Mo, Ha, elk, and sheep (Ov) PrP. **B**) Circular dichroism spectra of recombinant PrPs. **C**) Aggregation curves for rBVPrP (black), rMoPrP (red), rHaPrP (green), rElkPrP (brown), and rOvPrP (blue) as determined using a ThT fluorescence assay. **D**) Quantification of the lag phases for the ThT aggregation assays. **E**) Quantification of plateau ThT fluorescence values for the aggregation assays. **F**) Quantification of max ThT fluorescence:plateau ThT fluorescence ratios for the aggregation assays. For panels C-F, data are mean ± s.e.m. from 15 independent replicates. For panels D-F, statistical significance was assessed using one-way ANOVA with Dunnett’s multiple comparisons test. P-values for pairwise comparisons relative to rBVPrP are shown.

Next, we decided to test the effects of the key residues identified in the cultured cell prion infection experiments described above on the aggregation kinetics of rBvPrP. We purified rBVPrP with the MoPrP-equivalent residues at positions 155 and 170 as well as MoPrP with the BVPrP-equivalent residues at positions 154 and 169 (**Fig. 8A**). Both mutants were correctly refolded into an α-helical state (**Fig. 8B**). Remarkably, we found that rMoPrP with asparagine at residues 154 and 169 behaved identically to rBVPrP in the ThT aggregation assay, with a similarly reduced lag phase, enhanced maximum ThT fluorescence, and increased ratio of maximum:plateau ThT fluorescence (**Fig. 8C-F**). Conversely, rBVPrP lacking the two asparagine residues behaved similarly to rMoPrP. Finally, to examine the influence of C-terminal BVPrP residues on the aggregation kinetics of rPrP, we purified rBVPrP with four different residue combinations at positions 227 and 230 (**Fig. 9A**). All four rPrPs were successfully refolded into an α-helical state (**Fig. 9B**). As expected, rBVPrP aggregated rapidly in the ThT assay (**Fig. 9C**). However, rBVPrP containing the C-terminal residues at positions 227 and 230 from either mouse/hamster, human, or elk/ovine/dog/rabbit/bovine PrP aggregated even more rapidly with significantly shorter lag phases than wild-type rBVPrP (**Fig. 9C, D**). Therefore, there is a strong correlation between how rapidly a given rPrP spontaneously polymerizes into aggregates *in vitro* and the ability of the corresponding PrP^C^ to permit cross-species prion replication when expressed in cultured cells.

**Figure 8.**
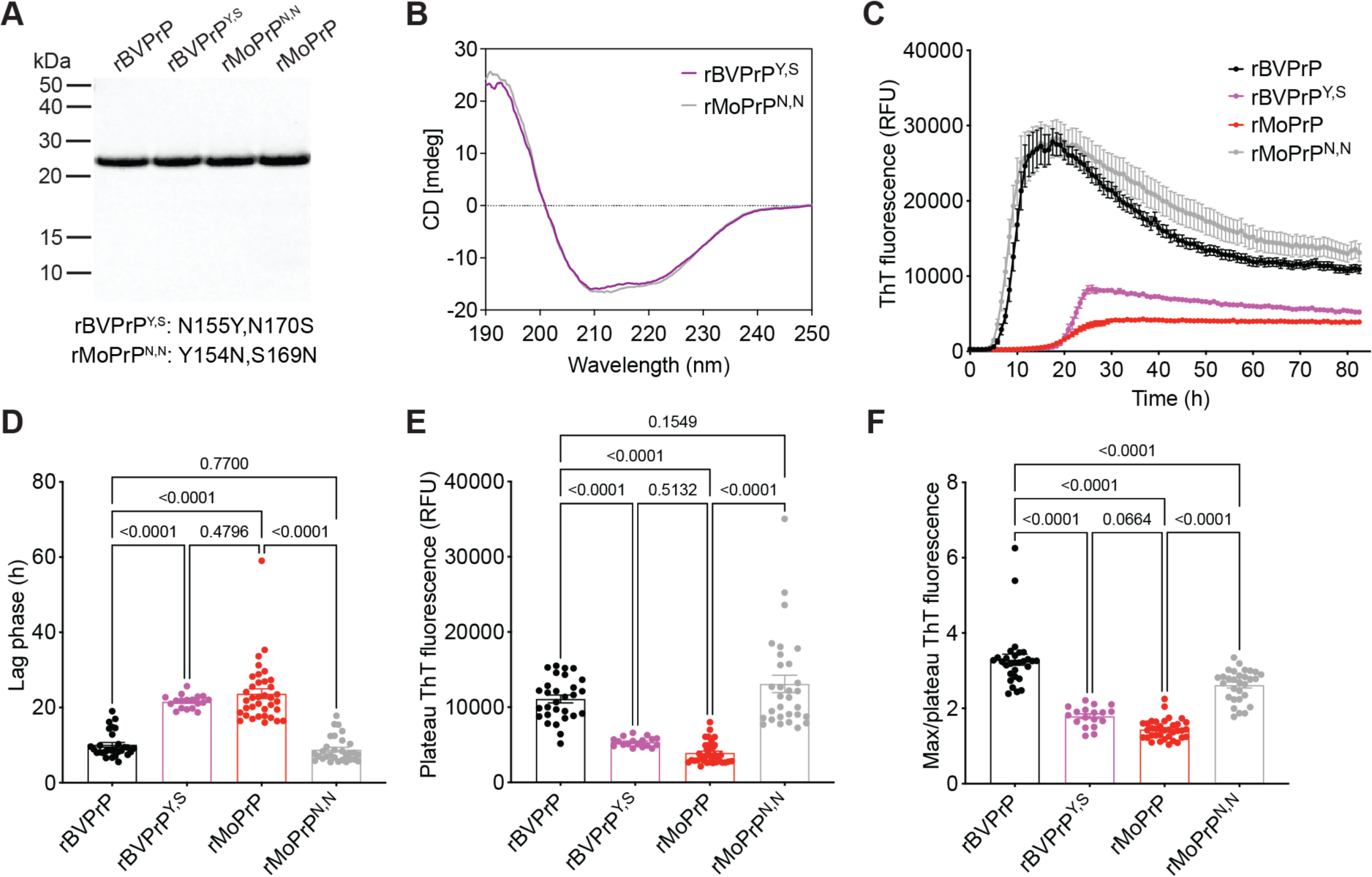
Asparagine residues at positions 155 and 170 of BVPrP promote accelerated spontaneous aggregation kinetics. **A**) SDS-PAGE followed by Coomassie blue staining of the indicated purified recombinant PrPs. **B**) Circular dichroism spectra of recombinant PrPs. **C**) Aggregation curves for rBVPrP (black, n = 29), rMoPrP (red, n = 36), rBVPrP with the N155Y/N170S substitutions (purple, n = 18), and rMoPrP with the Y154N/S169N substitutions (grey, n = 30) as determined using a ThT fluorescence assay. **D**) Quantification of the lag phases for the ThT aggregation assays. **E**) Quantification of plateau ThT fluorescence values for the aggregation assays. **F**) Quantification of max ThT fluorescence:plateau ThT fluorescence ratios for the aggregation assays. For panels C-F, data are mean ± s.e.m. For panels D-F, statistical significance was assessed using one-way ANOVA with Tukey’s multiple comparisons test. P-values for selected pairwise comparisons are shown.

**Figure 9.**
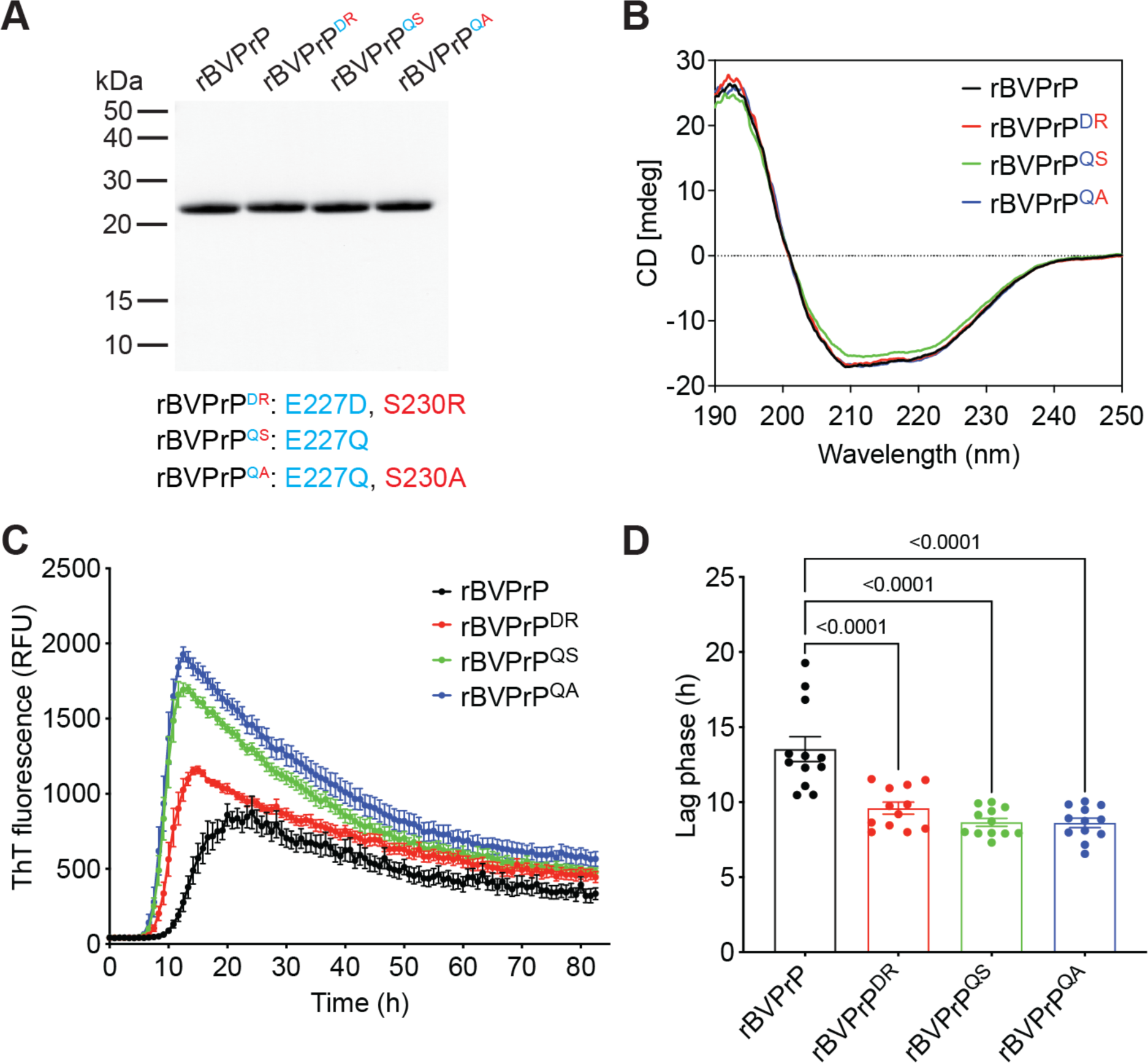
Residues E227 and S230 of BVPrP delay spontaneous aggregation. **A**) SDS-PAGE followed by Coomassie blue staining of the indicated purified recombinant PrPs. **B**) Circular dichroism spectra of recombinant PrPs. **C**) Aggregation curves for rBVPrP (black), rBVPrP with the E227D/S230R substitutions (red), rBVPrP with the E227Q substitution (green), and rBVPrP with the E227Q/S230A substitutions (blue) as determined using a ThT fluorescence assay. **D**) Quantification of the lag phases for the ThT aggregation assays. For panels C and D, data are mean ± s.e.m. from 12 independent replicates. For panel D, statistical significance was assessed using one-way ANOVA with Dunnett’s multiple comparisons test. P-values for pairwise comparisons relative to rBVPrP are shown.

## Discussion

The species barrier in prion disease is predominantly dictated by structural compatibility, which can be conferred by identical or highly similar amino acid sequences between PrP^Sc^ seed and PrP^C^ substrate (11). However, BVPrP demonstrates that a single PrP^C^ substrate can possess the appropriate residues to support the replication of most, if not all prion strains, despite the presence of sequence mismatches between PrP^C^ and PrP^Sc^. In this study, we leveraged the ability of BVPrP to permit cross-species prion replication when expressed in CAD5-PrP^-/-^ cells to identity the key regions in BVPrP responsible for the universal prion acceptor phenotype. Our findings suggest that BVPrP’s anomalous behavior is governed by five amino acid residues that are differentially present in the sequences of MoPrP, HaPrP, and BVPrP. The presence of asparagine residues at positions 155 and 170 of BVPrP explains why BVPrP can permit cross-species prion replication whereas MoPrP cannot. Although HaPrP contains these critical asparagine residues, it lacks essential BVPrP-specific residues at positions 112, 139, and 205, explaining why HaPrP is not a universal prion acceptor. Thus, in some ways, BVPrP can be considered a natural chimera between MoPrP and HaPrP that possesses the right combination of residues to enable cross-species prion replication. However, we unexpectedly found that C-terminal BVPrP-specific residues at positions 227 and 230 impede cross-species prion transmission, suggesting that the sequence of BVPrP may have been fine-tuned through evolution to counteract its intrinsic propensity to misfold. Furthermore, a striking correlation was also observed between the relative susceptibility of a given PrP variant to cross-species replication and the speed at which it forms aggregates spontaneously *in vitro*. Thus, the ability of PrP to adopt aggregation-competent structures may facilitate crossing of the species barrier.

The molecular details of how the five identified BVPrP residues enable cross-species prion replication remain unclear. Conceivably, the residues could act at the PrP^C^ level, potentially by rendering PrP^C^ more prone to aggregation due to localized structural changes that destabilize the protein or promote formation of β-sheet structures. Indeed, rBVPrP aggregated into β-sheet-rich structures much more rapidly than rPrPs from other species, indicative of a higher misfolding propensity and that rPrP, and therefore BVPrP^C^, may be intrinsically less stable than PrPs from other species. The structure of BVPrP^C^ is very similar to that of PrP^C^ from other mammalian species, except for a more “rigid loop” prior to the second α-helix that is partially mediated by residue N170, which is one of the key BVPrP residues we identified (56). The presence of a rigid loop has been shown to promote PrP misfolding and spontaneous prion formation as well as modulate interspecies prion transmission, although this appears to be driven by sequence rather than structural homology (57–62). When coupled with the observation that both elk and horse PrP^C^ also display a rigid loop but do not function as permissive substrates for cross-species prion replication (63, 64), this suggests that the rigid loop is not the major determinant of BVPrP’s atypical properties.

Alternatively, the key BVPrP-specific residues could act at the level of PrP^Sc^ or hypothetical misfolded intermediate PrP species that occur during the aggregation process. It has been speculated that the relative asparagine content in PrP is a good predictor of susceptibility to prion infection (35), and indeed we discovered that two asparagine residues in BVPrP (N155 and N170) are critical for cross-species prion replication. Due to their polar nature, asparagine side chains can stabilize β-sheets and promote the formation of highly stable zippers or polar clasps (65), potentially lowering energy barriers that would normally dissuade interspecies prion conversion. In the structure of mouse RML PrP^Sc^, residues V112 and I139 exist in close spatial proximity within the N-terminal lobe and are found within a hydrophobic pocket that is more tightly packed than in the corresponding region in hamster 263K PrP^Sc^ (5, 6). Thus, these residues may help to stabilize the structure of PrP^Sc^, enhancing the efficiency of cross-species prion conversion. In the structure of 263K PrP^Sc^, residue 205 is in proximity to residues near the C-terminal GPI anchor attachment site whereas in the structures of several mouse PrP^Sc^ strains it is distal to this region due to a distinct orientation of the PrP^Sc^ C-terminus (5–8, 66). Therefore, residue 205 could conceivably influence cross-species prion replication via structural modulation of the PrP C-terminal region. It is possible that the five identified BVPrP residues act via diverse and synergistic mechanisms involving both PrP^C^ destabilization and PrP^Sc^ stabilization to engender BVPrP with the ability to replicate many different prion strains.

A key finding of this study is the discovery of residues in the C-terminal region of BVPrP that impede cross-species prion replication. This was not simply due to a requirement for sequence identity at residues 227/230 for efficient prion replication, as insertion of a single glutamine residue at position 227 of BVPrP, which is found in the sequence of human PrP but absent in both MoPrP and HaPrP, rendered cells more susceptible to both mouse and hamster prion strains. Thus, despite its ability to function as a universal acceptor of prions, BVPrP can conceivably be rendered even more permissive to prion replication by modifying the amino acids located proximal to the GPI anchor attachment site. Residue E227 is the only truly unique residue in BVPrP as S230 is also found in the sequence of human PrP. It is tempting to speculate that the glutamic acid residue at position 227 arose due to evolutionary pressure to counteract the inherent misfolding propensity of BVPrP, potentially reducing the occurrence of spontaneous or transmitted prion disease in bank voles. Our findings are opposite to those of previous reports that found that BVPrP residues E227 and S230 are pivotal to the universal prion acceptor phenotype, collectively functioning as a linchpin domain to enable templated or spontaneous structural conversion (36, 37). The reasons for this discrepancy are unclear but could be related to the experimental paradigms employed. We utilized an *in vivo* cellular environment in which BVPrP^C^ is correctly attached to the outer leaflet of the plasma membrane and would be subject to normal cellular proteostasis mechanisms, whereas *in vitro* prion amplification assays employing a purified, non-membrane-anchored PrP^C^ substrate were used elsewhere (36). When the C-terminal region of PrP^C^ is not constrained via attachment to the cell membrane and/or sonication is used to fragment growing PrP^Sc^ aggregates, it is possible that the inhibition conferred by BVPrP residues 227/230 can be overridden. Nonetheless, our results further support a model in which the right combination of amino acids at positions 227/230 are important modulators of prion replication efficiency. Consistent with this notion, a mutation in the C-terminal region of PrP can modulate the kinetics of human prion replication (67). Exactly how residues 227/230 affect prion conversion remains to be determined but could be related to localized structural changes that alter PrP^C^ stability or affect interactions with components of the plasma membrane. One study found that the GPI anchor in human PrP is attached to residue G229, which means that residue S230 would not be present in the mature processed protein (68). Although, to the best of our knowledge, the precise attachment site for the GPI anchor in BVPrP has not been experimentally confirmed, this suggests that residue 227 may be a more critical determinant of cross-species prion replication, which is in line with our results.

The striking correlation between the relative permissiveness of a given PrP^C^ to replicate both hamster and mouse prions in cultured cells and the aggregation kinetics of the corresponding rPrP implies that there may be mechanistic links between cross-species prion replication and spontaneous prion generation. Indeed, in transgenic mice, expression of the I109 variant of BVPrP promotes both spontaneous prion formation and cross-species prion replication (23, 28). We speculate that BVPrP is more conformationally malleable than PrPs from other species, potentially allowing it to form misfolded intermediate species that can either further assemble to initiate spontaneous prion formation or be templated by pre-existing PrP^Sc^ in a manner that does not require precise sequence identity between PrP^C^ and PrP^Sc^. Whether C-terminal residues in BVPrP attenuate spontaneous prion generation remains to be determined, but we note that changing a single residue in BVPrP to its human equivalent (E227Q) significantly accelerated the spontaneous aggregation of rPrP. Although human PrP contains a misfolding-promoting residue at its C-terminus, it is unlikely to be prone to spontaneous prion conversion since it only possesses two of the five BVPrP residues we have shown to be critical for cross-species prion conversion (I139 and M205). This could explain why spontaneous prion diseases are very rare in humans.

There are several limitations to our study. First, we only used mouse and hamster prion strains to investigate BVPrP-mediated cross-species prion transmission. It is therefore unknown whether the same amino acid determinants in BVPrP govern cross-species transmission involving prion strains from sheep, cervids, cattle, or humans, all of which can be transmitted to bank voles and transgenic mice expressing BVPrP (16–18, 23). While both mouse and hamster prions result in robust PrP^res^ generation following infection of BVPrP-expressing CAD5-PrP^-/-^ cells, detection of infection in cells treated with chronic wasting disease prions is more challenging and requires amplification using RT-QuIC, which implies low levels of PrP^Sc^ accumulation (43, 44). Second, we only utilized a single readout of prion infection, relative levels of PrP^res^, to assess the ability of a given PrP to permit cross-species prion replication. Finally, although we have documented a strong parallel between susceptibility of PrP to cross-species prion replication and enhanced spontaneous aggregation kinetics of the corresponding rPrP, it must be noted that rPrP aggregates adopt fibrillar structures distinct from authentic PrP^Sc^ and lack the levels of infectivity displayed by brain-derived prions (69, 70). Furthermore, rPrP lacks the N-glycans that are present in cell-expressed PrP, and we found that a BVPrP mutant unable to undergo N-glycosylation was not susceptible to cross-species prion replication. Thus, further studies are required to fully understand any potential links between rPrP aggregation and cross-species prion replication.

## Acknowledgements

We are grateful to Stanley Prusiner for providing the HuM-D13 antibody.

## Declarations

### Availability of data and material

All data generated or analyzed during this study are included in this published article.

### Author contributions

HA, GSU, and JCW contributed to the study conception and design. Material preparation, data collection and analysis were performed by HA, ZP, ZAMA, LL, GA, SM, and SE. The first draft of the manuscript was written by HA and JCW, and all authors commented on previous versions of the manuscript. All authors read and approved the final manuscript.

