## Supplemental data for "The Molecular Determinants of a Universal Prion Acceptor"

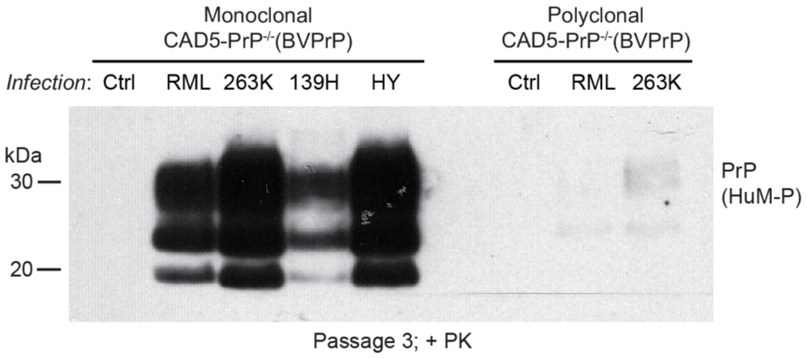


**Supplementary Figure 1. Infection of monoclonal and polyclonal CAD5-PrP$^{-/-}$ cell lines expressing BVPrP with mouse and hamster prions.** Immunoblot of PrP$^{res}$ levels in monoclonal and polyclonal lines of stably transfected CAD5-PrP$^{-/-}$(BVPrP) cells infected with either mouse (RML) or hamster (263K, 139H, or HY) prion strains. Cells were analyzed at passage 3 post-infection. PrP$^{res}$ was detected using the antibody HuM-P. Molecular weight markers indicate kDa.

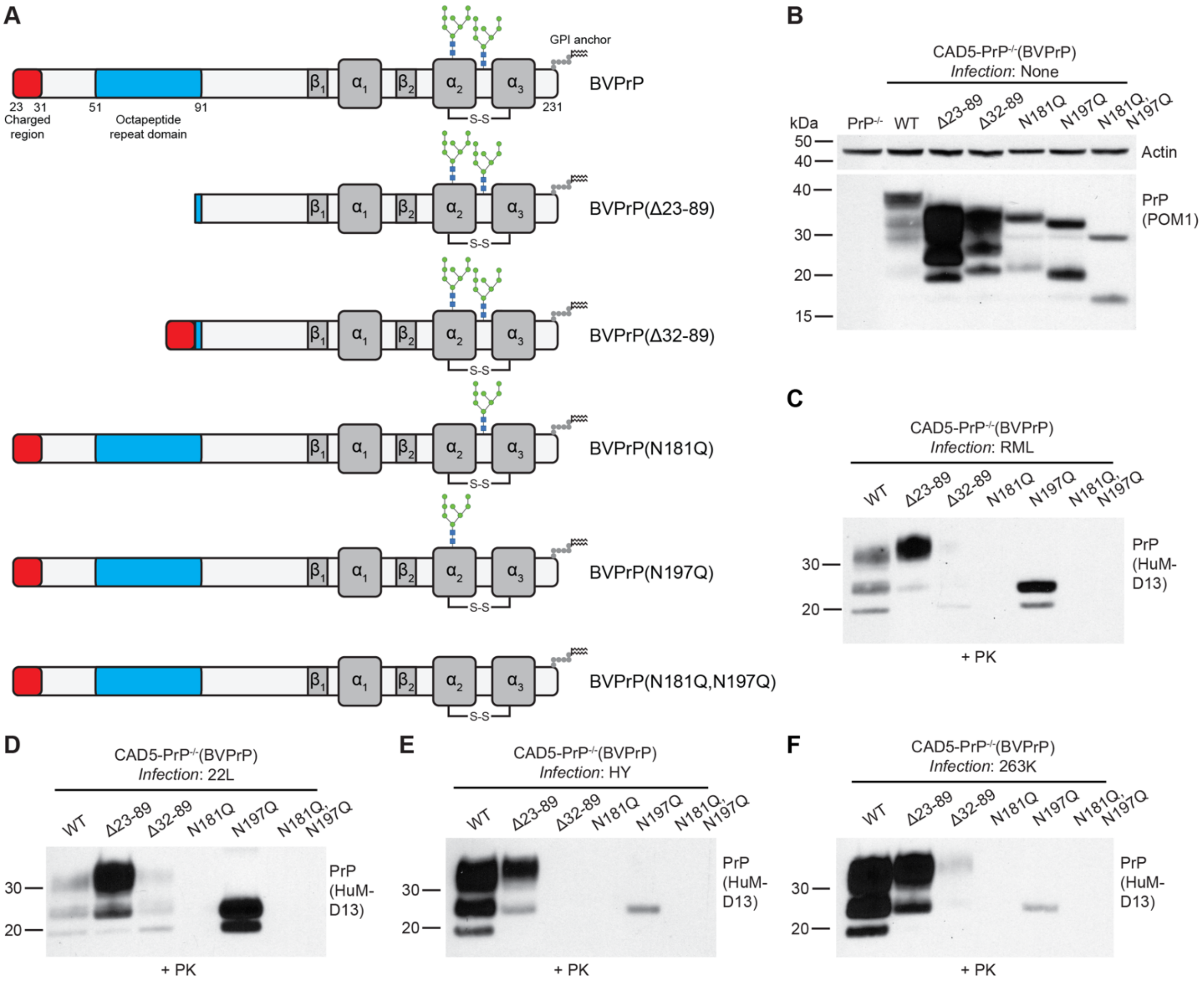


**Supplementary Figure 2. The role of N-glycosylation and the BVPrP N-terminal domain in cross-species prion transmission. A**) Domain structures of the BVPrP constructs used. For simplicity, the N- and C-terminal signal sequences are not depicted in the diagram. All constructs contain methionine at codon 109. **B**) Immunoblot for $PrP^C$ in undigested lysates from CAD5-$PrP^{-/-}$ cells stably expressing the indicated BVPrP constructs. The blot was reprobed with an antibody against actin. **C-F**) Immunoblots of $PrP^{res}$ levels in CAD5-$PrP^{-/-}$ cells stably expressing the indicated BVPrP constructs challenged with either mouse RML (**C**), mouse 22L (**D**), hamster HY (**E**), or hamster 263K (**F**) prions. $PrP^C$ was detected using the antibody POM1 whereas $PrP^{res}$ was detected using HuM-D13. In all panels, the molecular weight markers indicate kDa.

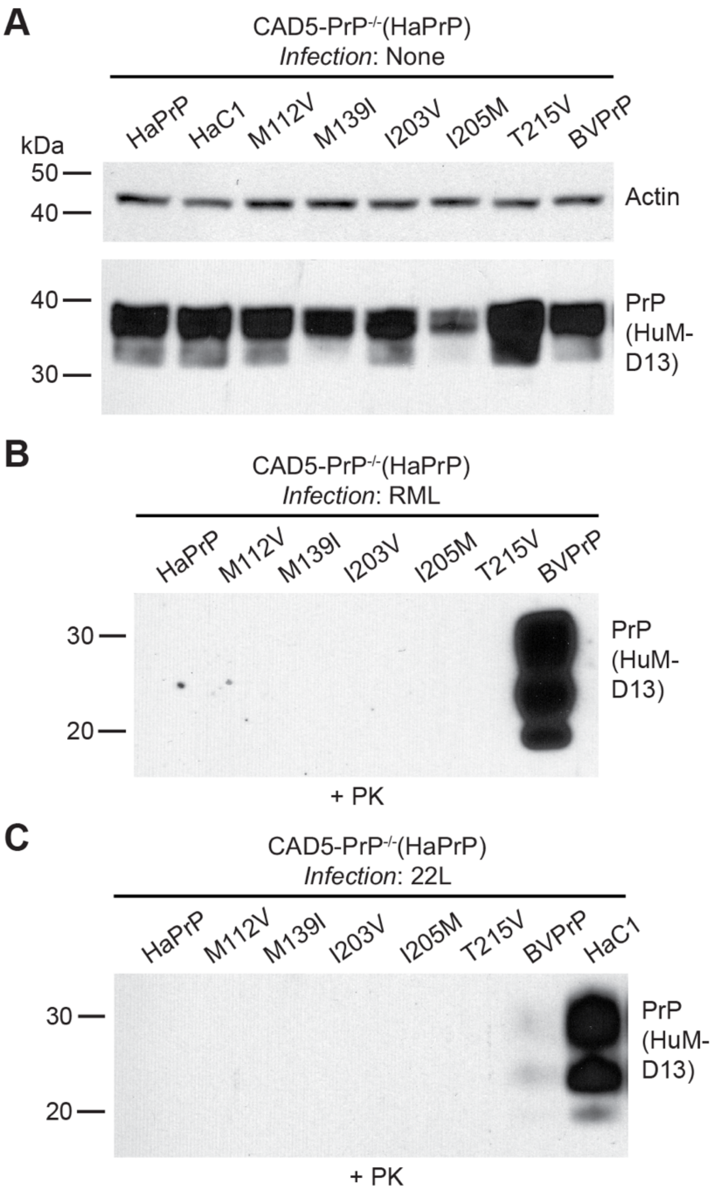


**Supplementary Figure 3. Single BVPrP residue substitutions in HaPrP are insufficient for enabling cross-species prion replication. A**) Immunoblot for $PrP^{C}$ in undigested lysates from CAD5-$PrP^{-/-}$ cells stably expressing wild-type HaPrP or BVPrP, the HaC1 chimera, or HaPrP with the indicated BVPrP residue substitutions. The blot was reprobed with an antibody against actin. **B-C**) Immunoblots of $PrP^{res}$ levels in lysates from CAD5-$PrP^{-/-}$ cells stably expressing the indicated PrPs challenged with either RML (**B**) or 22L (**C**) mouse prions. PrP detected using the antibody HuM-D13. In all panels, the molecular weight markers indicate kDa.
